# Learning the DNA syntax of human microbiomes to infer health and disease

**DOI:** 10.1101/2025.09.30.679483

**Authors:** Ana Mejia-Fleisacher, Noa Bossel Ben-Moshe, Tomer Antman, David Zeevi, Roi Avraham

## Abstract

The human microbiome is a key factor in human health and alterations in community structure are associated with diverse pathological conditions. However, defining universal criteria to distinguish healthy from altered microbiome configurations remains challenging due to inter- and intra-individual variability, database-dependent approaches, and the complexity of analyzing numerous microbial features simultaneously. Here, we developed an approach that learns the syntax of the entire DNA of human microbial communities, using Sequence-Informed GC-normalized 4-mers (SIG-mers) that feed into statistical and machine learning frameworks. We identified distinct SIG-mer signatures that differentiate microbiomes of body sites across diverse healthy human populations. These signatures reveal both global microbiome shifts and individual-specific dynamics in response to antibiotic treatments and in chronic inflammatory disease. Leveraging machine learning models, we inferred health- and disease-associated microbiome states from SIG-mer profiles, capturing the degree of perturbation and disease severity. Our findings highlight SIG-mer profiling as a robust, unbiased and broadly applicable approach for personalized microbiome diagnostics and guiding therapeutic interventions.

## Introduction

The human microbiome plays a pivotal role in maintaining health and modulating disease. The microbial communities present at defined niches across the human body are important drivers of host metabolism, immunity, and overall host physiology^1–3^. While the microbiome of healthy individuals generally maintains a dynamic equilibrium, perturbations can shift communities toward altered states that are detrimental to the host^4,5^. Deviations from healthy state are implicated in various medical conditions, including autoimmune^6^ and inflammatory diseases^7,8^, cancer^9,10^, psychiatric disorders^11^, cardiovascular disorders^12^, obesity^13,14^, and type 2 diabetes^15^. These findings have set the stage for defining microbiome-based biomarkers and therapeutic interventions^16^, highlighting the critical need to accurately distinguish healthy from altered states across individuals and populations. While these studies have been instrumental in identifying microbiome species that are associated with health and disease, establishing a universal consensus of ‘healthy’ microbiome characteristics remains challenging^17,18^.

Microbiome complexity and variability pose fundamental obstacles for establishing universal health signatures. Taxonomic approaches reveal substantial inter-individual variation in microbial species composition among healthy individuals^19–22^, with additional intra-individual shifts in some species abundance showing up to 100-fold fluctuations over short periods^23–26^. Geographic location, diet, and other environmental factors influence taxonomic profiles, further complicating efforts to identify stable compositional patterns^27–29^. Other efforts describing distinct enterotypes, stable community states defined by dominant bacterial genera that would transcend individual variation^30^, were complemented by subsequent taxonomic analyses, demonstrating that enterotypes represent points along a continuous spectrum rather than discrete categories^31,32^. Furthermore, large-scale genomic studies continue to expand the known repertoire of microbial species and genomes, revealing vast taxonomic diversity that remains uncharacterized^20,33–36^.

Beyond these complexities, annotation-dependent approaches rely on existing reference databases, which restricts analyses to previously characterized microorganisms and introduces bias toward highly characterized taxa^37^. Consequently, a fraction of sequencing reads, particularly from understudied body regions, remains unmapped and discarded, reducing community profile resolution and overlooking yet uncharacterized bacteria as well as key members such as fungi and viruses^38,39^. Additionally, alignment methods suffer from misclassification of reads from closely related species, poor detection of low-abundance taxa, and an inability to accurately resolve strain-level differences that may drive phenotypic variation. De novo assembly-based methods offer a reference-free alternative, addressing some of these limitations by generating metagenome-assembled genomes (MAGs) from previously uncharacterized microbial populations^40^. However, these methods require extensive sequencing depth, and their reliability has been questioned due to frequent contamination, incompleteness, presence of chimeric species populations, and uneven recovery of population-variable genes^35,41–43^.

An alternative strategy is an unbiased analysis of the DNA sequence of the whole community, bypassing the need for genome- or gene-centric comparisons. Specifically, early studies applied k-mer analysis, such as 4-mer frequencies, to describe genomic signatures, indicating they are conserved within species, distinct between taxa, and informative for detecting phylogenetic relationships^44–46^. Yet, most studies using k-mer analysis have focused on comparative genomics, including taxonomic profiling, genome assembly, and phylogenetic inference. More recently, 4-mer frequency patterns in environmental metagenomes were shown to describe the selective pressures that shape microbial communities, driven by differences in community GC content stemming from the biome of origin^47,48^.

In this study, we applied 4-mer analysis to human microbiomes and observed significant GC biases between different body sites. To directly survey the DNA syntax independent of these biases, we developed a refined 4-mer metric by sequence-informed GC normalized features (SIG-mers). Leveraging metagenomic data from healthy individuals across diverse geographic locations and datasets, we identify unique SIG-mer patterns that characterize stable, organ-specific microbiomes. We then explored how these profiles are altered during acute antibiotic perturbations and in chronic diseases. Using regression-based machine learning models, we inferred alterations in microbiome composition from SIG-mer profiles and detected both population-wide community shifts associated with disease severity, and individual-specific microbiome dynamics. We propose that our approach will provide robust and predictive analytics as a microbiome assessment framework, complementary to reference-based analyses.

## Results

### GC normalized 4-mers captured metagenomic fingerprints of healthy human organs

To globally analyze DNA signatures of healthy human microbiomes, we applied 4-mers frequency analysis to infer profiles of microbial communities across different body sites (**Fig. 1A**). This DNA sequence-driven method describes each community’s metagenome using a standardized set of 136 unique features present across genomes and samples. We analyzed 19 publicly available metagenomic datasets encompassing 3,349 samples from 1,482 healthy individuals across three organs and eight anatomical sites, using 1M reads per sample to remove sequencing depth biases (**Table 1**). Hierarchical clustering of 4-mers profiles revealed four clusters driven by GC-content of 4-mers (**fig. S1A**), reflecting differential GC-content between the metagenomes of different body sites (*p* < 0.001, *F*(7, 3341) = 727, ANOVA with Tukey post-hoc test, **fig. S1B, Table 2**). To extract sequence-informed 4-mers independent of GC content, we normalized the observed 4-mers frequencies by their expected frequencies based on each sample’s GC content, creating Sequence-Informed GC-normalized 4-mers (SIG-mers; see Methods, **Fig. 1A**). Clustering SIG-mer scores, we observed no GC-driven clusters (**fig. S1C, Table 3**). To validate our normalization, we compared 4-mer frequencies and SIG-mer scores, stratified by the number of GC’s in each 4-mer across samples with different overall GC compositions (**fig. S1D, E**). This analysis indicated that before normalization, low-GC samples were enriched for AT-rich 4-mers (e.g., AAAA) while depleted for GC-rich 4-mers (e.g., GCGC), with the opposite pattern observed in high-GC samples (**fig. S1D**). This GC bias was eliminated after normalization, as demonstrated by SIG-mer scores (**fig. S1E**). Next, we evaluated pairwise SIG-mer score differences across Hamming distances, confirming that 4-mers with similar sequences are not systematically more similar in their scores than those with divergent sequences (*p* > 0.05, Kruskal–Wallis, **fig. S1F**). Together, these analyses indicate that SIG-mers capture sequence-informed patterns independent of constraints such as GC content of samples and sequence similarity.

**Fig. 1.**
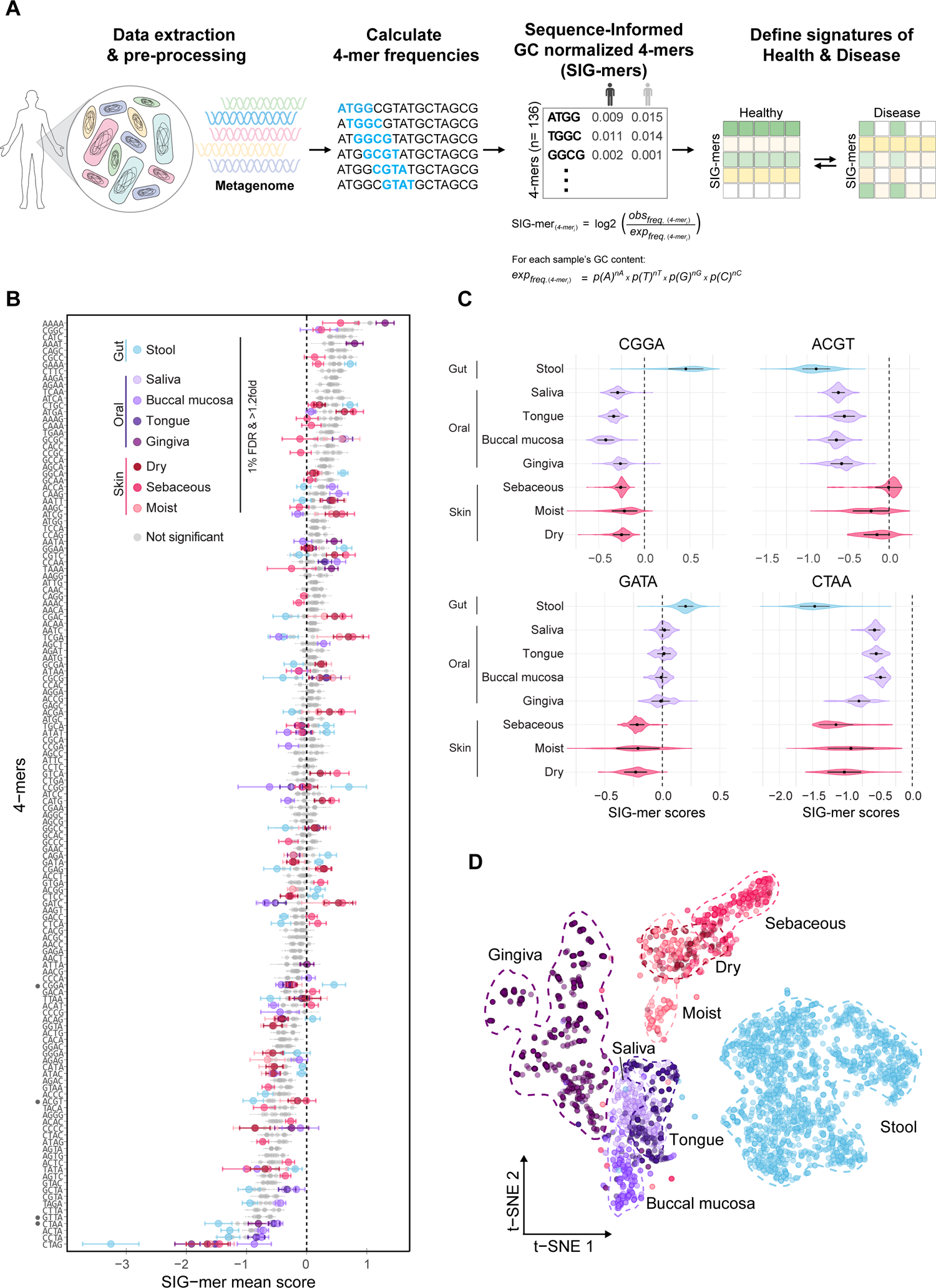
GC normalized 4-mers capture metagenomic fingerprints of healthy human organs. **A.** Pipeline overview. Shotgun metagenomic reads undergo preprocessing and quality control, followed by the calculation of 4-mer frequencies using a sliding window approach. Frequencies are GC-normalized to provide SIG-mer profiles. **B.** Differential abundance analysis of SIG-mers (two-sample *t*-test, 1% FDR, >1.2-fold) comparing each site against all others. The dotplot shows the mean SIG-mer score for each 4-mer at each site. Significant 4-mers are colored according to site, while non-significant 4-mers are shown in grey. Data included: Gut (stool, n = 1530); Oral (saliva, n = 189; buccal mucosa, n = 170; tongue, n = 239; gingiva, n = 650); Skin (dry, n = 93; sebaceous, n = 273; moist, n = 205). **C.** Violin plots showing SIG-mer scores across samples for 4-mers that were significant in the differential abundance analysis (panel B), illustrating their distribution across sites. **D.** t-SNE visualization of SIG-mer profiles from the same healthy samples shown in panel B. Each point represents an individual sample, colored according to the anatomical site of origin.

To study SIG-mer patterns across different body sites, we plotted the mean SIG-mer score for each 4-mer at each site. Specific SIG-mers showed global directional biases, deviating from zero consistently across sites. For example, CTAG (bottom 4-mer, **Fig. 1B**) was consistently underrepresented in all sites, particularly in gut samples. CTAG has been independently reported as a markedly rare sequence in microbial genomes, marking a conserved site for restriction-modification systems and genome regulation^49–52^. Similarly, AAAA (Top 4-mer, **Fig. 1B**) was overrepresented at all sites, consistent with its high genomic frequency due to base composition bias and DNA structural roles^53–55^. We next compared each site against all others and found distinct SIG-mers that displayed organ- and site-specific deviations (indicated by colored dots, two-sample *t*-test, 1% FDR & >1.2-fold, **Fig. 1B**, **Table 4**). Notably, several SIG-mers had opposite directional shifts between organs but consistent biases within sites of the same organ. For example, we plotted SIG-mer scores for each sample and observed that organ- and site-specific deviations are evident also beyond inter-individual variation (**Fig. 1C**). CGGA was enriched in gut sites but underrepresented in other organs, while GATA was overrepresented in gut and underrepresented across all skin sites. Even when deviations were directionally similar across organs, such as for ACGT and CTAA, the magnitude of deviation differed significantly between sites, revealing fine-scale, site-specific microbial patterns. Thus, beyond the inter-individual variability, SIG-mer analysis captures site-specific patterns of healthy microbiomes.

To test whether these SIG-mer patterns can be reproduced at lower sequencing depths, we performed similar analyses using lower read coverage (10K and 100K reads) and compared the results to our original analysis (1M reads). Correlation analyses showed high stability of SIG-mer profiles across sequencing depths, with significant correlations between 1M and 100K coverage (*Pearson* r = 0.998) and between 1M and 10K coverage (*Pearson* r = 0.996) (**fig. S1G**).

Principal component analysis (PCA) of SIG-mers profiles indicated a clear separation between healthy gut samples and other organs, supported by significant differences between sites (ANOSIM: R = 0.835, *p* = 0.001; PERMANOVA: R² = 0.647, *p* = 0.001) (**fig. S1H**).

Dimensionality reduction using *t*-distributed stochastic neighbor embedding (*t*-SNE) demonstrated similar separation between organs and partial separation of anatomical sites within the organs (**Fig. 1D**). Gut samples could be further stratified by geographic origin, consistent with previously documented regional differences of gut microbiomes^27–29^ (**fig. S1I**). Finally, we compared SIG-mers of mouse gut metagenomics data^56^ to healthy human organs, and measured higher correlation between mouse gut and human gut (*Pearson* r = 0.95) than correlations between the mouse gut and other human body sites (**fig. S1J**), indicating conservation of SIG-mer profiles across species within the same organ.

Overall, we demonstrate that SIG-mers can robustly define organ and site-specific metagenomic patterns across species, enabling consistent characterization of healthy microbiome communities across individuals, body sites, geography and datasets.

### SIG-mers captured dynamic changes of the gut microbiome in different antibiotic regimens

Antibiotics are known to cause collateral damage to the gut microbiome, leading to acute changes in diversity, composition, and function^57–59^. While loss of microbial diversity and recovery after antibiotic treatment are well-documented, inter-individual differences in baseline microbiome composition limit the ability to quantify the dynamics of changes, treatment-specific effects and recovery rates across individuals.

To evaluate antibiotic-induced perturbations, we applied our SIG-mer approach to mouse metagenomic data comprising mouse stool samples collected before and during seven different antibiotic treatments^56^ (**Table 1**). We compared treated (72 hours) and naïve samples across all treatments to identify changes in SIG-mer profiles (two-sample *t*-test, 1% FDR & >1.2-fold; **Fig. 2A, fig. S2A, Table 5**, **Table 6**). These analyses revealed global dose-dependent shifts in SIG-mer profiles during antibiotic treatment relative to baseline. Only a few SIG-mers were significantly changed under low-dose, single-antibiotic regimes (fosfo-low: 4; cipro-low: 2), whereas a large number of significant changes of SIG-mers were observed under high-dose antibiotic cocktails (amp+cipro-low: 8; amp+cipro+fosfo-low: 38; amp+fosfo-high: 75; cipro+fosfo-high: 71). Consistent with these differential analyses, correlation analysis identified two distinct clusters: one comprising naïve and low-dose single treatments, and another comprising cocktail antibiotic treatments, reflecting the observed shifts in SIG-mer profiles (**fig. S2B**).

**Fig. 2.**
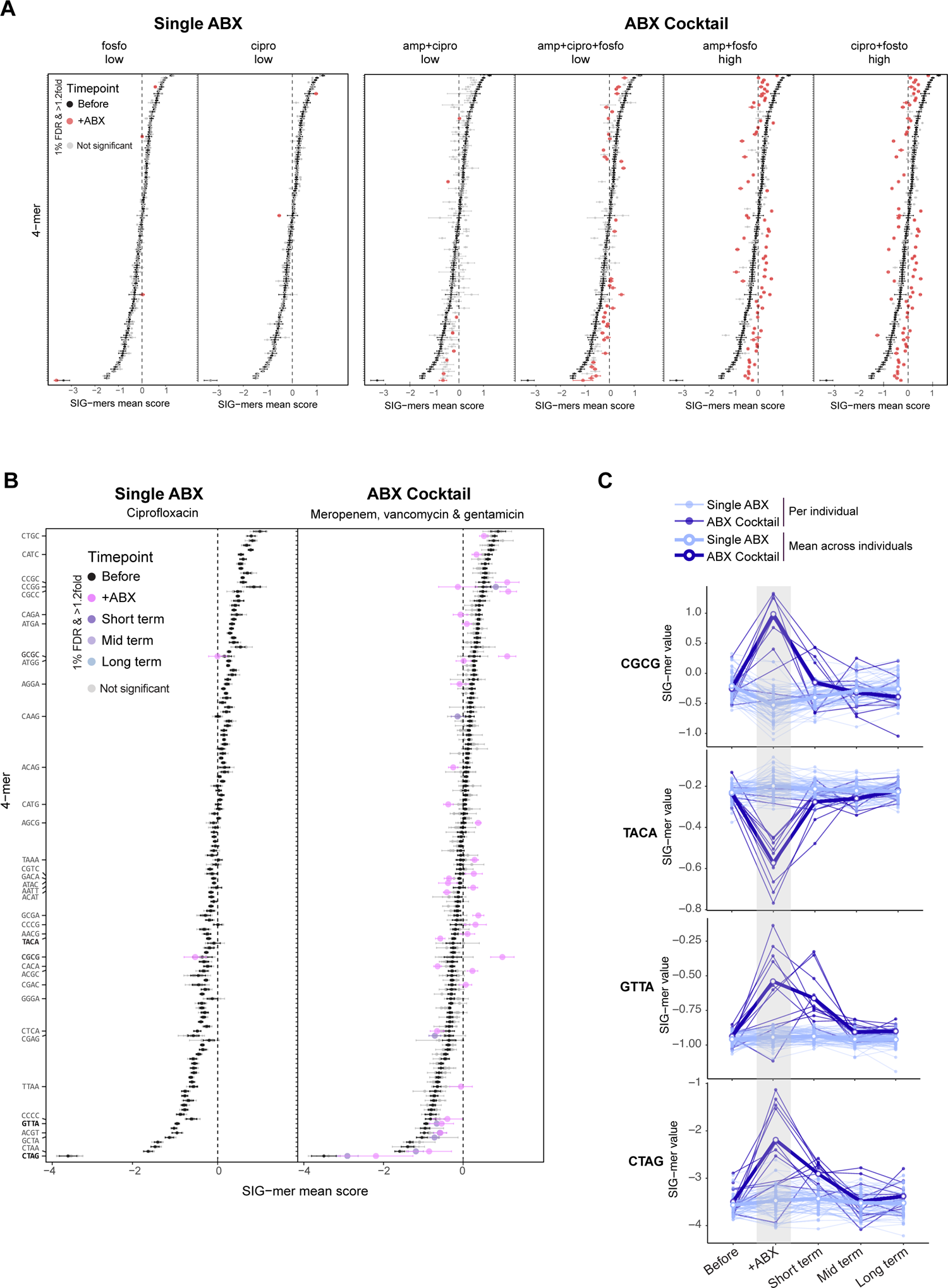
SIG-mers captured dynamic changes of the gut microbiome in different antibiotic regimens. A. Mouse longitudinal antibiotic treatment dataset. **A.** Differential abundance analysis of SIG-mers (two-sample *t*-test, 1% FDR, >1.2-fold) comparing treated samples at 72 hours post-treatment with naïve mice. Each dotplot represents one antibiotic regimen, from low-dose single-antibiotic to high-dose cocktail treatments, showing the mean SIG-mer score for each 4-mer in naïve mice (black dots) and treated mice. Significant 4-mers are colored in red, while non-significant 4-mers are shown in grey. **B-C. Human longitudinal antibiotic treatment datasets. B.** Differential abundance analysis of SIG-mers (two-sample *t*-test, 1% FDR, >1.2-fold) comparing baseline (before treatment) samples with samples collected at multiple timepoints across individuals: during treatment (day 5 for single-antibiotic, day 4 for cocktail), short-term (day 8), mid-term (day 28 for single-antibiotic, day 42 for cocktail), and long-term (day 77 for single-antibiotic, day 180 for cocktail). Dotplots show mean SIG-mer scores per dataset, with the left plot for single-antibiotic treatment and the right plot for cocktail treatment. Significant 4-mers are colored by timepoint, and non-significant 4-mers are shown in grey. **C.** Individual dynamics of SIG-mer scores over time for selected 4-mers (same timepoints as in panel B) in both datasets: single-antibiotic treatment (light blue) and cocktail treatment (dark blue). Mean trajectories across individuals per dataset are shown with thicker lines and white dots.

To determine SIG-mer profile changes under antibiotic treatments in human microbiomes, we analyzed longitudinal metagenomic data from two studies of human stool samples collected before, following antibiotic treatment, and during long-term recovery of healthy individuals^60,61^ (**Table 1**). The first dataset^61^ included 60 healthy volunteers who received ciprofloxacin twice daily for 5 days and were followed for 20 weeks. We observed subtle changes in SIG-mer profiles, with only two SIG-mers significantly changed between baseline and day 5 (two-sample t-test, 1% FDR & >1.2-fold; **Fig. 2B**, **fig. S2C, Table 7**, **Table 8**). The second dataset^60^ included 12 healthy volunteers who received a daily antibiotic cocktail (meropenem, vancomycin, and gentamicin) for 4 days, with follow-up over 180 days. Here, we observed several SIG-mers that were significantly changed between baseline and after 4 days treatment (two-sample *t*-test, 1% FDR & >1.2-fold; **Fig. 2B, fig. S2D, Table 9**, **Table 10**). To capture individual-level dynamics underlying the observed changes, we analyzed longitudinal SIG-mer trajectories for each individual across both datasets. We observed that certain SIG-mers display different dynamics: CGCG changed significantly in both datasets but in opposite directions and magnitudes; TACA showed a similar bidirectional trend but did not reach significance in the ciprofloxacin cohort.; GTTA and CTAG remained stable under single-antibiotic treatment yet shifted significantly under cocktail treatment (**Fig. 2C**). Additionally, CTAG and GTTA trajectories highlighted heterogeneous responses and recovery patterns among individuals, varying in both timing and magnitude. PCA indicated similar patterns, with clear separation between baseline (day 0), antibiotic-cocktail and single-antibiotic regimens, with the two treatments shifting in distinct directions (**fig. S2E**).

Overall, SIG-mer analysis robustly identifies acute gut microbiome perturbations, capturing both different individual dynamics and dose-dependent changes across treatment regimens beyond inter-individual variation.

### SIG-mer-based prediction model distinguished mild and severe antibiotic perturbation dynamics

Having established that SIG-mers can robustly describe gut microbial population compositions in health and antibiotic-perturbed microbiomes, we next assessed the predictive power of our approach. We developed a LASSO-regularized logistic regression model ^62,63^ to classify microbiome samples as healthy or perturbed, based on their SIG-mer scores. The model was trained and validated on samples from both antibiotic treatment datasets, using pre-treatment samples as “healthy” (day 0) and samples from final day of treatment as “perturbed” (day 5 for single antibiotics, day 4 for cocktails) (**fig. S2C, S2D**). The model outputs a probability score between 0 and 1, where values closer to 1 indicate a higher likelihood of microbiome perturbation. We assessed the model’s performance using the prediction probabilities from leave-one-out cross-validation (LOOCV), where in each round one individual’s samples were withheld for testing while the model was trained on all remaining data^64^ (**fig. S3A**). The model achieved a predictive performance with an area under the receiver operating characteristic curve (AUC) of 0.90 (**Fig. 3A**), 89% accuracy, and area under the precision-recall curve (AUPRC) of 0.91 (**fig. S3B**). Using this model, we could discriminate between healthy and perturbed microbiomes in the two datasets. We then leveraged the variable coefficients from LASSO iterations to identify the most contributing features for classifications (see Methods) and used them to train a final model. The resulting model generates a dysbiosis risk score based on a refined set of 34 SIG-mers (**fig. S3C**).

**Fig. 3.**
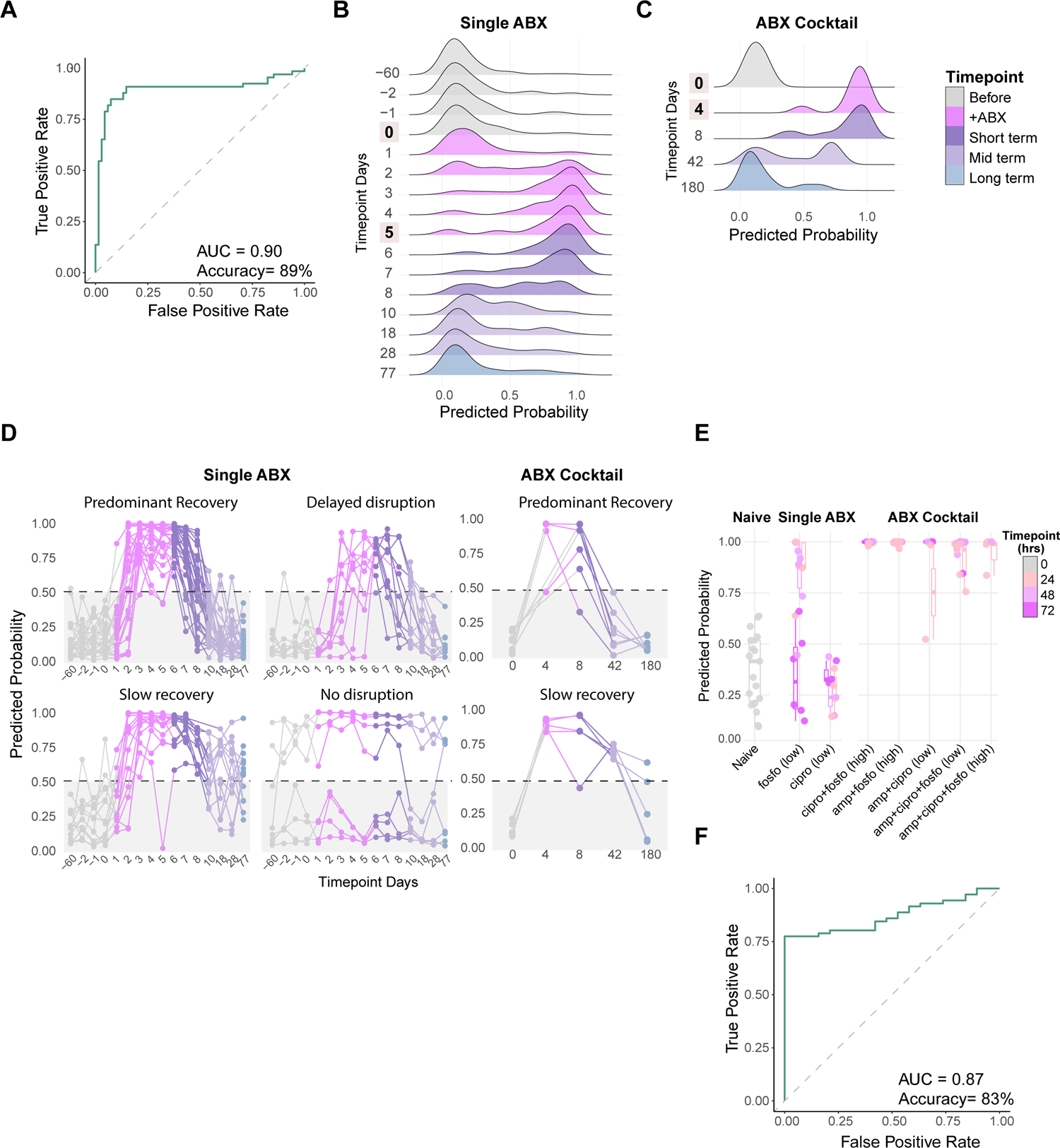
SIG-mer-based prediction model distinguishes mild and severe antibiotic perturbation dynamics. **A.** ROC curve showing model performance based on LOOCV predicted probabilities for healthy versus antibiotic-treated samples. **B-C.** Density plots of predicted probabilities for all samples by timepoint. Training timepoints (day 0 and final treatment day) are indicated; remaining timepoints represent unseen test data. **B.** Single-antibiotic dataset. **C.** Antibiotic-cocktail dataset. **D.** Individual recovery dynamics categorized based on predicted probabilities from the model. Dashed line indicates default classification threshold at 0.5 for both antibiotic datasets. **E.** Model performance on an independent mouse dataset. Boxplot shows predicted probabilities for samples at each timepoint across treatment groups. **F.** ROC curve showing model performance for naïve versus treated mouse samples.

To study microbiome dynamics during antibiotic treatments, we applied the final model to infer microbiome status on all remaining timepoints that were excluded from the training stage. In the single antibiotic treatment dataset, three additional pre-treatment timepoints (days −60, −2, and −1) were correctly predicted as healthy with no significant difference from day 0 (*p* > 0.05; Wilcoxon test, Bonferroni correction; **Table 11**). Antibiotic-mediated disruption of SIG-mer profiles was observed beginning at day 2 (*p* < 0.001; Wilcoxon test, Bonferroni correction; **Table 11, 3B**) and peaking at days 3-4 during treatment. Recovery dynamics back to a predicted healthy state started around days 7-8, returning to baseline levels, with no significant differences between day 0 and day 18 (*p* > 0.05; Wilcoxon test, Bonferroni correction; **Table 11**) (**Fig. 3B**). Similar predicted dynamics were observed for the cocktail antibiotic treatment (**Fig. 3C**), with significant changes relative to baseline for day 4 and 8 (*p* < 0.001; Wilcoxon test, Bonferroni correction; **Table 12**) and in some individuals, changes seemed to persist through the mid-term timepoint (day 42), but without reaching significance (*p* > 0.05; Wilcoxon test, Bonferroni correction).

To examine differences in individual dynamics, we set a threshold of 0.5 on the model probability scores, to distinguish healthy from disrupted states, and identified several recurring patterns of recovery dynamics (**Fig. 3D**). The predominant recovery trajectory, consistent with previously reported patterns^59,65^, was characterized by early disruption during treatment (day 2 for single-ABX, day 4 for ABX-cocktail), followed by a return to healthy composition several days after treatment cessation (day 10 for single-ABX and day 42 for cocktail). Alternatively, we observed different patterns of recovery: (1) slow or incomplete recovery; recovery was evident only at day 77 for single-ABX or day 180 for ABX-cocktail, while in some cases perturbation was observed until the last time point measured; (2) Delayed disruption; delayed disruption with rapid baseline return (for single-ABX only). (3) Non-disrupted individuals fall into two groups: those predicted as healthy throughout all timepoints, and those predicted as disrupted from baseline. This latter group may reflect underlying individual differences in baseline microbiome compositions that limit our model predictions. These results indicate that our SIG-mer approach and prediction model can distinguish between individuals that are more susceptible to microbiome perturbations and ones more resilient, laying the foundations for an in depth understanding of microbiome vulnerabilities in the population.

To evaluate model accuracy and to estimate prediction at reduced sequencing coverage, we subsampled the datasets to 100K and 10K reads per sample and repeated the full analysis pipeline: SIG-mer calculation followed by LASSO-regularized logistic regression, training on two time points and testing on all remaining timepoints. Model performance remained consistent across all coverage levels, with an AUC of 0.9 and AUPRC of 0.91 for 100K reads, and an AUC of 0.9 and AUPRC of 0.89 for 10K reads, with comparable classifications, compared to 1M reads (**fig. S3D**).

Finally, we tested the model on an independent dataset using the mouse antibiotics experiment (**Fig. 2A**). The model, trained on human datasets, predicted mouse gut alterations for all ABX-cocktail treatments compared to baseline (**Fig. 3E,** *p* < 0.01 vs. naive; Wilcoxon test, Bonferroni correction; **Table 13**), but not all single antibiotic treatments were predicted as perturbed. Model performance was measured with an AUC of 0.87, an accuracy of 83%, and an AUPRC of 0.97 (**Fig. 3F, fig. S3E**). We further examined an additional human dataset of perturbed gut microbiomes from 26 intensive care unit (ICU) patients (Anthony et al., 2022). This dataset represents a population with high antibiotic exposure and extensive medical interventions. We compared baseline samples from the antibiotic datasets (day 0**, fig. S2C, S2D**) to the ICU patients and measured significant changes in SIG-mer profiles (two-sample *t*-test, 1 % FDR, **fig. S3F, S3G, Table 14**). We then applied our model and measured significant differences in predicted probabilities between ICU patients and baseline samples (**fig. S3H**, *p* <0.001, Wilcoxon test).

Together, these results demonstrate that our SIG-mer-based model provides robust prediction of microbiome alterations and recovery dynamics in the gut microbiome, beyond inter-individual variation and applicable across datasets and organisms. Model validation on independent datasets and different organism demonstrates that the human-trained model captures robust and conserved features of antibiotic-induced gut changes.

### SIG-mer-based classification model predicted chronic disease

Chronic inflammatory diseases are marked by recurrent episodes^66,67^ and considerable heterogeneity in disease manifestation^68^, often associated with altered microbiome composition^2,69^. We next set out to test whether our predictive models can discriminate disease states beyond patient heterogeneity.

Atopic dermatitis (AD) is a common chronic inflammatory skin disorder, actively influenced by skin microbes that modulate host immunity and exacerbate lesions^70–72^. We analyzed an AD dataset^73^, including 29 children with AD and 8 healthy controls (**Table 1**). Metagenomics data was collected from skin swabs in moist sites at distinct time points: baseline (stable disease state), flare (skin disease exacerbation without recent therapies), and post-flare (10–14 days after initiating skin-directed treatment). In each visit, disease severity was scored dividing patients into mild, moderate, and severe categories^74^. We performed SIG-mer analysis on the data and stratified differential abundance across disease severity categories relative to healthy controls (**fig. S4A, Table 15**). We mainly observed changes in SIG-mer profiles of patients with severe disease, with additional alterations in mild and moderate categories (two-sample *t*-test, 1% FDR & >1.2-fold, **Fig. 4A**, **Table 16**). Pearson correlation analysis indicated a progressive divergence in microbiome profiles from healthy to severe states (**fig. S4B**). PCA analysis indicated significant changes between severe samples and other groups (ANOSIM: R = 0.24, p = 0.001; PERMANOVA, R² = 0.22, p = 0.001) (**fig. S4C**).

**Fig. 4.**
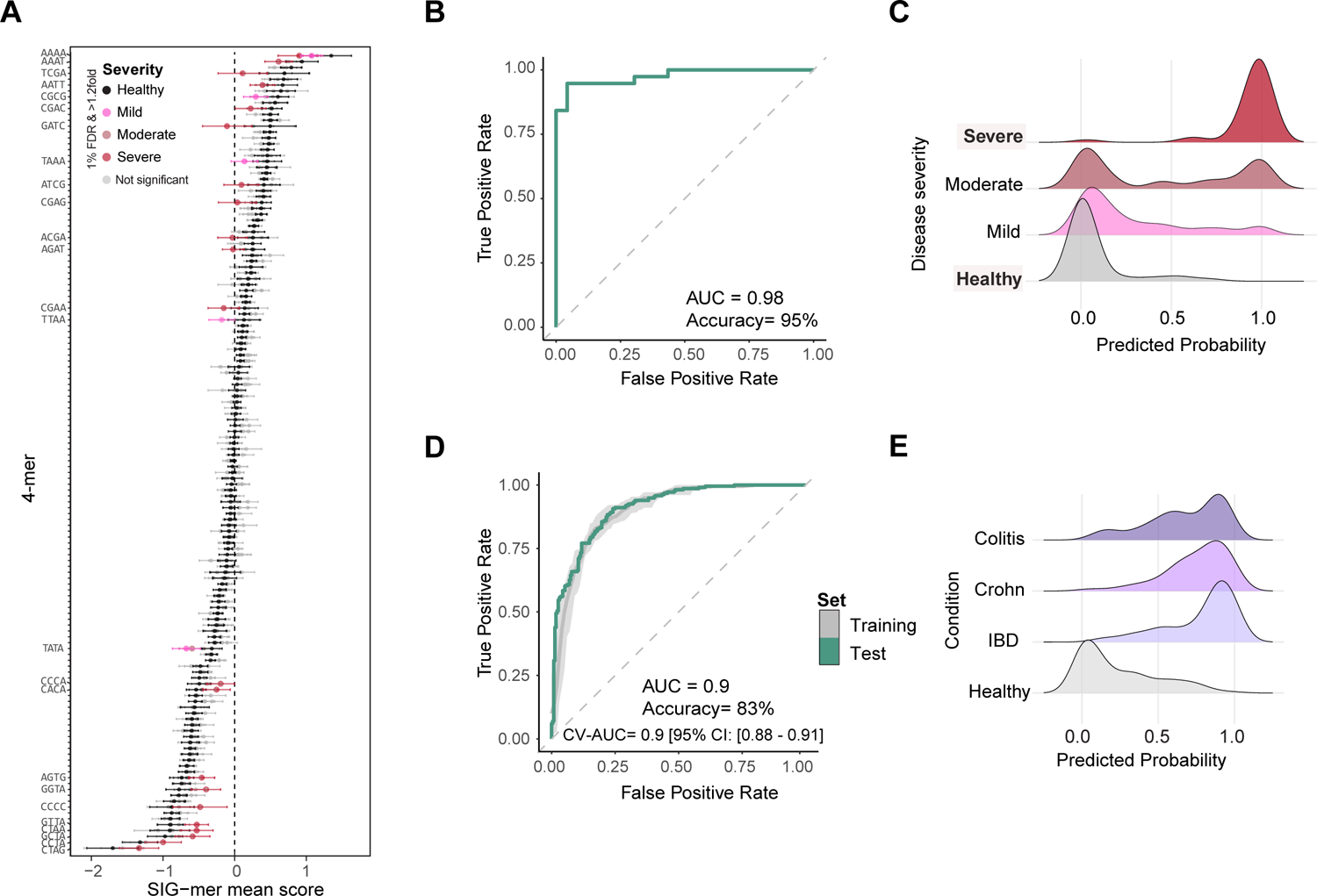
SIG-mer-based classification model predicts chronic disease. A-C. Atopic dermatitis dataset. **A.** Differential abundance analysis of SIG-mers (two-sample *t*-test, 1% FDR, >1.2-fold) comparing healthy samples with atopic dermatitis samples across different disease severity categories. Dot plot shows the mean SIG-mer score for each 4-mer in healthy samples (black) and in each severity category. Significant 4-mers are colored by severity category, while non-significant 4-mers are shown in grey. **B.** ROC curve showing model performance based on LOOCV predicted probabilities for healthy versus severe samples. **C.** Density plots of predicted probabilities for all samples stratified by disease severity category. **D-F. IBD datasets. D.** ROC curve showing model performance for healthy versus IBD subtypes. Grey: 5-fold cross-validation; green: independent test set. **E.** Density plots of test set predicted probabilities by IBD subtype. Significant differences observed between healthy and all IBD categories distributions.

Next, we trained a LASSO-regularized logistic regression model, employing LOOCV to distinguish between severe AD and healthy configurations. In each iteration, all samples from one individual were excluded from training and used for validation. The model robustly classified severe disease samples with clear separation from healthy samples (**fig. S4D**), with an AUC of 0.98 (**Fig. 4B**), an accuracy of 0.95%, and an AUPRC of 0.99 (**fig. S4E**). We then trained a final model, utilizing 27 SIG-mers with highest mean coefficients (**fig. S4F**). We applied the model to infer the status of remaining samples (**Fig. 4C**). Moderate and mild distributions were significantly different from healthy and severe (*p* < 0.001; Wilcoxon test, Bonferroni correction; **Table 17**). Mild samples were mostly classified as healthy (77% of samples), while in the moderate category, samples categorized as severe were from the post-flare timepoint (100% of samples) (**fig. S4G**).

Next, we examined SIG-mer profiles in datasets collected from gut-associated chronic inflammatory bowel disease patients (IBD). IBD are characterized by chronic relapsing inflammation of the digestive tract, with two major forms: Crohn’s disease (CD) affecting discontinuous intestinal regions, and ulcerative colitis (UC) restricted to the colon^75,76^. Gut microbiota alterations are considered both a clinical symptom and causative driver of disease progression^77,78^. To evaluate the utility of our method for IBD prediction, we analyzed 6 datasets comprising 936 samples: healthy controls (n =229), general IBD (n = 190), Crohn’s disease (n = 316), and ulcerative colitis (n = 201) (**Table 1**). We performed SIG-mer analysis and observed no significant changes between healthy and disease states (two-sample *t*-test, 1% FDR & >1.2-fold, **fig. S4H, fig. S4I, Table 18**, **Table 19**). Closer examination indicated significant but minor (<1.2-fold) changes in few SIG-mer scores relative to healthy microbiome composition (**fig. S4J)**.

We hypothesized that a classification model could capture subtle SIG-mer changes in disease states. We trained a LASSO-regularized logistic regression model to distinguish healthy individuals from those with any subtype of IBD. The model was trained on a randomly selected 70% of samples and evaluated using 5-fold cross-validation stratified by class and dataset. Based on feature coefficients from cross-validation, 102 SIG-mers were selected to build the final model (**Table 20**). Evaluation on the independent test set (30% of samples) demonstrated robust performance, with an AUC of 0.9 and accuracy of 83% (**Fig. 4D, fig. S4K**), as well as a PR-AUPRC of 0.9 (**fig. S4L**). Prediction probabilities significantly differed between IBD subtypes and healthy controls (*p* < 0.001; Wilcoxon test, Bonferroni correction; **Table 21**), with consistent performance across all IBD subtypes (**Fig. 4E**). Stratifying by dataset did not reveal any dataset-specific biases (**fig. S4M**).

Thus, our prediction models based on SIG-mer scores can robustly discriminate between healthy and inflammatory disease patients, providing an approach to classify microbiomes across disease, patients and datasets.

## Discussion

In this study we introduce SIG-mers, a reference-free approach that analyzes the DNA syntax of microbiomes. Using this approach, we discerned organ- and site-specific patterns detected in metagenomic data from healthy individuals. We propose that these profiles likely reflect local environmental constraints that select for well-adapted and similar microbial communities (**Fig. 1B**). The similarity between human and mouse gut SIG-mer profiles points to possible evolutionary conservation of sequence-level constraints in mammalian gut environments (**fig. S1J**). SIG-mers analysis captured broad ecological distinctions, such as between body sites, as well as finer-scale patterns like geographic origins of samples (**Fig. 1D, fig. S1I**). These patterns can also be reliably detected even at very low sequencing depths (as few as 10K reads (**Fig. S1G**)), where taxonomic approaches typically lose resolution^79,80^. These findings demonstrate that, as in environmental microbiomes^47,48^, the human microbiome retains clear sequence-level imprints of its surrounding physiological conditions. Together with the consistency of SIG-mer profiles across individuals and geographic locations, we propose that SIG-mers may reflect the conserved core of microbiome functional repertoires^19,81^, dominated by essential housekeeping and central metabolic pathways, and represent a promising focus for future studies.

SIG-mer profiles consist of a fixed set of features shared across genomes and samples, providing a standardized representation of microbial patterns. This standardization facilitated the utility of SIG-mers in logistic regression models as a powerful tool to distinguish healthy microbiomes from altered states (**Fig. 3A, 4B** and **4D**). Indeed, our model trained on antibiotic treatment data not only reproduced documented recovery trajectories^60,61,82,83^, but also captured finer inter-individual recovery dynamics (**Fig. 3D**). Furthermore, the model identified stable baseline configurations within individuals, with all pre-treatment timepoints classified as healthy (**Fig. 3B**), despite documented short-term temporal fluctuations in species abundance^23–26^. We propose that our approach provides a framework for studying community-wide behaviors, that may allow classifications of resilient from susceptible communities that could underlie disease states. By capturing the full genomic content of a metagenomic sample, SIG-mer analysis complements current approaches that are limited by overrepresentation of well-characterized organisms^37^, noisy variations driven by less abundant taxa, or underrepresentation of previously uncharacterized taxa, including protists, viruses, and fungi^38,39^. Moreover, the ability of our human-trained model to successfully classify mouse antibiotic treatment data (**Fig. 3E, 3F**) suggests that SIG-mer profiling captures general microbiome health principles extending beyond host-specific microbial communities. While SIG-mer analysis currently provides limited functional interpretability, future studies may uncover mechanistic insight by focusing on specific genomic regions, decomposing taxon-level contributions, and linking SIG-mer signals to gene-level functional profiles. Furthermore, as the number of samples increases, it may be possible to use longer k-mers, expanding feature space and thereby refining functional relevance, and signal resolution of our approach^84–86^.

Our analyses of disease-associated microbiome alterations in chronic inflammatory diseases could distinguish between milder and more severe states in AD patients based on SIG-mer profiles of the skin microbiome. Taxonomy-based microbiome classifications in this disease describe a highly heterogeneous disease state, with uneven reductions or enrichments of species, taxa, and functional groups between individuals. This suggests that at least part of the observed heterogeneity may stem from taxon-level variability, where community-level signals like SIG-mers could help to mitigate. In IBD, studies have documented altered gut microbial composition and metabolic pathways, enabling microbiome-based models to predict disease state^87^. Using our approach, we achieved comparable performance but still observed substantial variability between individuals within different disease subtypes. Future integration of larger datasets and clinical information on disease activity could further refine SIG-mer–based models, enabling the investigation of individual disease dynamics and responses to treatment. Achieving this will require within-individual sampling across different disease states, which could support the development of more personalized diagnostic and predictive tools.

## Methods and materials

### Meta-analysis of publicly available metagenomic datasets

We downloaded publicly available human and mouse shotgun metagenomic samples from three major repositories: the European Nucleotide Archive (ENA), the Sequence Read Archive (SRA), and the National Genomics Data Center (CNCB). In total, we collected 7,087 metagenomes from 22 different studies (**Table 1**). These samples span nine countries, with the most substantial contributions from the United States (n = 5922), China (n = 408), Denmark (n= 206), and South Africa (n=164). Samples were collected from three major body sites: sites: gut/stool (n = 3,407), oral cavity (n = 1,843), and skin (n = 1,837).

Inclusion criteria for healthy individuals varied across studies but generally excluded participants who had received antibiotic treatment within the preceding six months or had conditions known to be associated with microbiome alterations.

### Mouse dataset

Shotgun metagenomic fastq files were obtained from SRA (study accession number: PRJNA886985; Table 1). The study used 8–10-week-old female Balb/c mice treated with different combinations of three antibiotics: ampicillin (amp), ciprofloxacin (cipro), and fosfomycin (fosfo). Treatment groups included: triple combination, dual combinations, and monotherapies, administered at high and low concentrations. High-dose treatments were 200 mg/kg ampicillin, 50 mg/kg ciprofloxacin (both twice daily), and 1000 mg/kg fosfomycin (once daily). All treatments lasted 3 days. Stool samples were collected from the distal ileum and proximal colon at 24, 48, and 72 hours post-treatment initiation.

### Single antibiotics

Shotgun metagenomic fastq files were obtained from SRA (study accession number: PRJNA974858; Table 1). The study analyzed fecal samples from 60 healthy adults living in the USA who received a 5-day treatment of twice-daily ciprofloxacin (500 mg) administered orally. Samples were collected at multiple timepoints, which we categorized as: before (9 weeks before and 3 days immediately before treatment), +ABX (daily during days 1–5), short-term (days 6–8), mid-term (days 10, 18, 28), and long-term (day 77).

### Cocktail antibiotics

Shotgun metagenomic fastq files were obtained from SRA (study accession number: PRJEB20800; Table 1). The study analyzed fecal samples from 12 healthy Caucasian males who received a 4-day treatment of once-daily antibiotic cocktail consisting of 500 mg meropenem, 500 mg vancomycin, and 40 mg gentamicin dissolved in apple juice and administered orally. Samples were collected at five timepoints, which we categorized as: before (day 0, pre-antibiotic), +ABX (day 4, last day of treatment), short-term (day 8), mid-term (day 42), and long-term (day 180).

### ICU dataset

Shotgun metagenomic fastq files were obtained from SRA (study accession number: PRJNA703034; Table 1). The ICU fecal specimens were convenience samples of remnant stool from the Clinical Microbiology Laboratory collected from 26 ICU inpatients who had diagnostic testing for Clostridioides difficile at Barnes-Jewish Hospital. No additional clinical metadata was available for these samples.

### Atopic dermatitis

The study analyzed skin samples from children with atopic dermatitis (AD) and healthy controls. After quality control, 29 AD patients and 8 healthy controls were retained for analysis. Skin swab samples were collected from sites of disease predilections, specifically Pc and Ac moist sites, during clinic visits. These visits were categorized as: stable disease or before flare (B), acute disease flare (F), and post-flare (PF) when inflammation had attenuated 10–14 days after initiating skin-directed treatment. Disease severity was assessed at each visit using the SCORAD assessment tool and categorized according to established conversion scales^74,88^: total SCORAD 0–24 (mild), 25–50 (moderate), and 51–103 (severe).

### Sequence data preprocessing

Downloaded fastq files underwent rigorous quality control to remove adapters, low-quality sequences, and host-derived DNA using the following sequential pipeline. We first used **SeqKit sana**^89^ with default parameters to sanitize corrupted single-line fastq files. Next, **Fastp**^90^ was applied for adapter removal, low-quality base trimming, poly-G tail trimming, and filtering out short reads using the following parameters: *--trim_front1 10, --trim_tail1 10, --length_required 50, --complexity_threshold 20, --trim_poly_g, --detect_adapter_for_pe, --low_complexity_filter and --overrepresentation_analysis*. **Dedupe** from BBMap suite (Bushnell, B., <u>BBMap</u>) was then used to remove duplicate reads with default parameters, followed by **SeqKit pair**^89^ to remove unpaired-end reads using default parameters. Finally, **Bowtie2**^91^ was used to thoroughly remove host-derived reads by aligning against the human telomere-to-telomere genome^92^ or mouse-telomere-to-telomere genome ^93^. Alignments were performed using the *--very-sensitive-local* parameter setting, and the mapped sequences were removed from subsequent analysis.

### 4-mer Frequency Calculation

For each sample, the first one million reads were used to count all possible tetranucleotide (4-mer) combinations independently for forward and reverse reads, using a sliding window of size 1. Reverse complementary sequences (e.g., “AAAA” and “TTTT”) were merged and treated as a single 4-mer (e.g., “AAAA/TTTT”), resulting in a 136-dimensional feature vector representing all unique 4-mer combinations per sample. After merging paired-end read count tables, raw 4-mer counts were normalized by dividing each value by the total number of 4-mers in the sample to obtain 4-mer frequencies. Variability in total 4-mer counts, despite using an equal number of reads, stems from differences in read lengths following adapter and low-quality base trimming ^47^. Additionally, the same procedure was used to calculate frequencies using reduced read numbers (10K and 100K reads) to evaluate the effect of sequencing depth on signature stability and to accommodate datasets where many samples fell below 1M reads after quality control. All analyses of the mouse dataset (**Fig. 2A, fig. S2A-B** and **Fig. 3E-F, fig. S3E**) and atopic dermatitis dataset (**Fig. 4A-C, fig. S4A-G**) were performed using frequencies calculated from 100K.

### Sequence-Informed GC-normalized 4-mers (SIG-mers)

4-mer frequencies are inherently biased by a sample’s GC content. To identify 4-mers that are over- or under-represented beyond this bias, frequencies were normalized based on each sample’s GC composition. For each sample, nucleotide probabilities were calculated as P(G) = P(C) = GC_content/2 and P(A) = P(T) = (1-GC_content)/2, where GC_content represents the sample’s overall GC composition. Expected 4-mer probabilities were then computed as the product of constituent nucleotide probabilities (e.g., P(ATGC) = P(A) × P(T) × P(G) × P(C)). The log₂ ratio of observed to expected frequencies was calculated: log₂(observed/expected) to quantify 4-mer enrichment or depletion. Complete SIG-mer data for each dataset are provided in the supplementary tables (**Tables 3, 5, 7, 9, 15,** and **18**).

### Differential abundance analysis

For each 4-mer, we performed a two-sample t-test comparing the condition of interest (e.g., sites, timepoints, disease severity categories, or IBD subtypes) against the corresponding healthy or baseline samples. Resulting p-values were adjusted for multiple testing using a 1% false discovery rate (FDR). To define significance, we required both an adjusted p-value below the threshold and an absolute fold-change greater than 1.2 (equivalent to a mean difference of ±0.263 on the log2 scale). Complete results, including raw p-values, adjusted p-values, and fold-change values for each analysis, are provided in the supplementary tables (**Table 4, 6, 8, 10, 16,** and **19**).

### Fold-change Heatmap visualization and clustering

For each sample, fold-change values were calculated relative to the appropriate reference: the paired baseline (before treatment) in longitudinal datasets or the mean of healthy controls in cross-sectional comparisons. Heatmaps display log₂ fold-change values of 4-mer scores, with rows representing 4-mers and columns representing individual samples. Samples were grouped by experimental category (e.g., treatment group, timepoint, or disease severity) to facilitate comparison of patterns across conditions.

### Classification models and cross-validation

#### General methodology

Linear classification regression models were performed using glmnet R package (version 4.1-8). All models employed LASSO regularization to remove features not contributing to prediction or that are redundant, with feature coefficients and lambda values recorded for each cross-validation fold. The mean absolute coefficient was calculated for each feature across all folds, and features with values greater than zero were selected. Final models were trained on the complete training dataset using the median lambda value from cross-validation with ridge regularization to retain all selected features. Model performance was evaluated using ROC curves to determine optimal classification thresholds and calculate accuracy, as well as precision-recall curves. Discriminative performance across categories was assessed by comparing predicted probability distributions using Wilcox test with BH correction.

#### Antibiotic model

Training data included baseline (day 0) and last-day of treatment samples (day 5 single-antibiotic, day 4 antibiotic-cocktail; n=120 single, n=24 cocktail). Leave-one-individual-out cross-validation was performed, resulting in 34 selected features. The final model was applied to all unseen remaining timepoint samples. To evaluate model robustness across sequencing depths, equivalent models were constructed using 100K and 10K read datasets. The 1M read model was further validated on two independent datasets (mouse and ICU datasets).

#### Atopic dermatitis model

Using healthy and severe samples (n=23 healthy, n=38 severe), this model followed the same cross-validation and feature selection procedure, yielding 27 selected features. The final model was applied to independent samples from mild and moderate disease severity categories.

#### IBD model

The dataset comprised 6 IBD datasets (UC, CD, unclassified IBD, and healthy) and 3 additional healthy datasets to address class imbalance (**Table 1**). Data were randomly split into training (70%, n=1,025) and test (30%, n=444) sets. The training set underwent 5-fold stratified cross-validation (stratified by disease status and dataset) for model development and feature selection, yielding 102 features (**Table 18**). Cross-validation performance on the training set is presented as AUC and PR-AUC curves (mean ± 95% CI). Final model performance was evaluated on the independent test set.

### Clustering and data visualization

All statistical analyses and data visualizations were performed in R version 4.4.1. Data visualization was conducted using ggplot2 (version 3.5.1) for general plotting, ggbeeswarm (version 0.7.2) for violin plots, ggridges (version 0.5.6) for ridge plots, and ggrepel (version 0.9.5) for annotations on the volcano plot. Dimensionality reduction was performed using Rtsne (version 0.17) for t-SNE, and density contours were overlaid using geom_density_2d() in ggplot2. To better represent density, contour lines were drawn with bins = 8 for low-density sites and bins = 40 for high-density sites, such as gingiva and stool. Heatmaps were generated using pheatmap (version 1.0.12) and ComplexHeatmap (version 2.20.0). Classification models were built using glmnet (version 4.1-8), and community ecology analyses were conducted using the vegan package (version 2.6-10) for ADONIS and ANOSIM tests.

